# Ataxin-2, Twenty-four and Dicer-2 are components of a non-canonical cytoplasmic polyadenylation complex

**DOI:** 10.1101/2021.12.17.472273

**Authors:** Hima Priyanka Nadimpalli, Tanit Guitart, Olga Coll, Fátima Gebauer

## Abstract

Cytoplasmic polyadenylation is a mechanism to promote mRNA translation in a wide variety of biological contexts. A canonical complex centered around the conserved RNA-binding protein family CPEB has been shown to be responsible for this process. We have previously reported evidence for an alternative non-canonical, CPEB-independent complex in *Drosophila*, of which the RNA-interference factor Dicer-2 is a component. Here, we investigate Dicer-2 mRNA targets and protein co-factors in cytoplasmic polyadenylation. Using RIP-Seq analysis we identify hundreds of novel Dicer-2 target transcripts, ∼50% of which were previously found as targets of the cytoplasmic poly(A) polymerase Wispy, suggesting widespread roles of Dicer-2 in cytoplasmic polyadenylation. Large-scale immunoprecipitation revealed Ataxin-2 and Twenty-four among the high-confidence interactors of Dicer-2. Functional analysis indicate that both factors form an RNA-independent complex with Dicer-2, and are required for cytoplasmic polyadenylation of Dicer-2 targets. Our results reveal the composition of a novel cytoplasmic polyadenylation complex that operates during *Drosophila* early embryogenesis.

## INTRODUCTION

Changes in poly(A) tail length chiefly contribute to the stability and translation of mRNA (Passmore and Coller, 2021). Poly(A) tail elongation has been shown to promote translation in biological contexts as diverse as oocyte maturation, early embryonic development, neuronal plasticity, bone formation, cell proliferation, senescence, inflammation, metabolism, circadian gene expression, hibernation and cancer (reviewed in D’Ambrogio et al., 2013; Ivshina et al., 2014; Kozlov et al., 2021; Kojima et al., 2012; Grabek et al., 2015; Gewartowska et al., 2021). This process is controlled by the cytoplasmic polyadenylation element binding (CPEB) family of proteins, which bind to U-rich cytoplasmic polyadenylation elements (CPEs) in the 3’ UTR of transcripts (Ivshina et al., 2014). Depending on sequence context and CPEB phosphorylation status, CPEBs can nucleate the formation of regulatory complexes that lead to either activation or repression of translation by balancing interactions of CPEB-associated factors with the mRNA cap and poly(A) tail (Piqué et al. 2008; Villalba et al., 2011; Fernández-Miranda and Mendez, 2012; Charlesworth et al., 2013; Ivshina et al., 2014). For example, phosphorylated CPEB1 recruits the cytoplasmic poly(A) polymerase Gld-2 to promote mRNA polyadenylation and translation, while unphosphorylated CPEB1 recruits silencing proteins that maintain the poly(A) tail short and the mRNA cap blocked.

CPEBs are highly conserved from humans to *Aplysia* (Ivshina et al., 2014). In *Drosophila*, the CPEB proteins Orb and Orb2 associate with the Gld-2 homolog Wispy to promote cytoplasmic polyadenylation (Cui et al., 2013; Norvell et al., 2015; reviewed in Kozlov et al., 2021). However, CPEB-independent cytoplasmic polyadenylation activities also exist in this organism (Coll et al., 2010; Dufourt et al., 2017). We have recently reported that the RNAi-related endoribonuclease Dicer-2 promotes cytoplasmic polyadenylation and translation of *Toll* mRNA in early embryos (Coll et al., 2018). Dicer-2 associates with structured elements in the 3’ UTR of *Toll* and interacts with Wispy to promote polyadenylation. The extent of Dicer-2-mediated polyadenylation and the factors that cooperate with Dicer-2 in this endeavor have remained uncharacterized. Here, we perform RNA-immunoprecipitation (RIP) and Dicer-2 pull-downs to identify mRNA targets and co-factors of Dicer-2 in cytoplasmic polyadenylation. We find that Dicer-2 binds to a large number of mRNAs that are also bound by Wispy, suggesting pervasive roles in cytoplasmic polyadenylation. We identify 86 high-confident Dicer-2 protein interactors, most of which are unrelated to RNA interference, suggesting broader functions of Dicer-2 in cell biology. Of these, 21 are RNA binding proteins, including the translation regulatory factors Twenty-four (Tyf) and Ataxin-2 (Atx2). We show that Tyf and Atx2 form an RNA-independent complex with Dicer-2 and Wispy, and are involved in cytoplasmic polyadenylation of Dicer-2 targets. These results reveal the composition of a non-canonical cytoplasmic polyadenylation complex, and point to an emerging diversity of polyadenylation regulators beyond CPEB.

## RESULTS AND DISCUSSION

### Identification of Dicer-2 mRNA targets

To identify the repertoire of RNAs that are bound by Dicer-2 in *Drosophila melanogaster*, we performed Dicer-2 immunoprecipitation (IP) from early (90 minutes) embryo extracts followed by RNA sequencing (RIP-Seq) in 6 biological replicates (see typical IP efficiency in Fig 1A). Parallel pull-downs with non-specific IgG were carried as negative controls. RNAs significantly associated to Dicer-2 were determined by comparison with matched inputs and with control IgG IPs (Figure 1B and Table S1). We considered RNAs present in the Dicer-2 IP in an excess larger than 1.5-fold in both comparisons, and with a Benjamini-adjusted p-value <0.001. This yielded a total of 1750 RNAs associated to Dicer-2, the vast majority of which (> 98 %) were mRNAs (Figure 1C). These results agree with the observation that, despite the central role of Dicer in small RNA biogenesis, most Dicer binding sites in human cells and *C. elegans* reside on mRNAs (Rybak-Wolf et al., 2014). These sites have been called ‘passive’ because their recognition by Dicer does not result in mRNA cleavage. On average, Dicer-2 mRNA targets contain a higher GC content indicative of increased structure compared to non-targets (Figure 1D).

**Figure 1.**
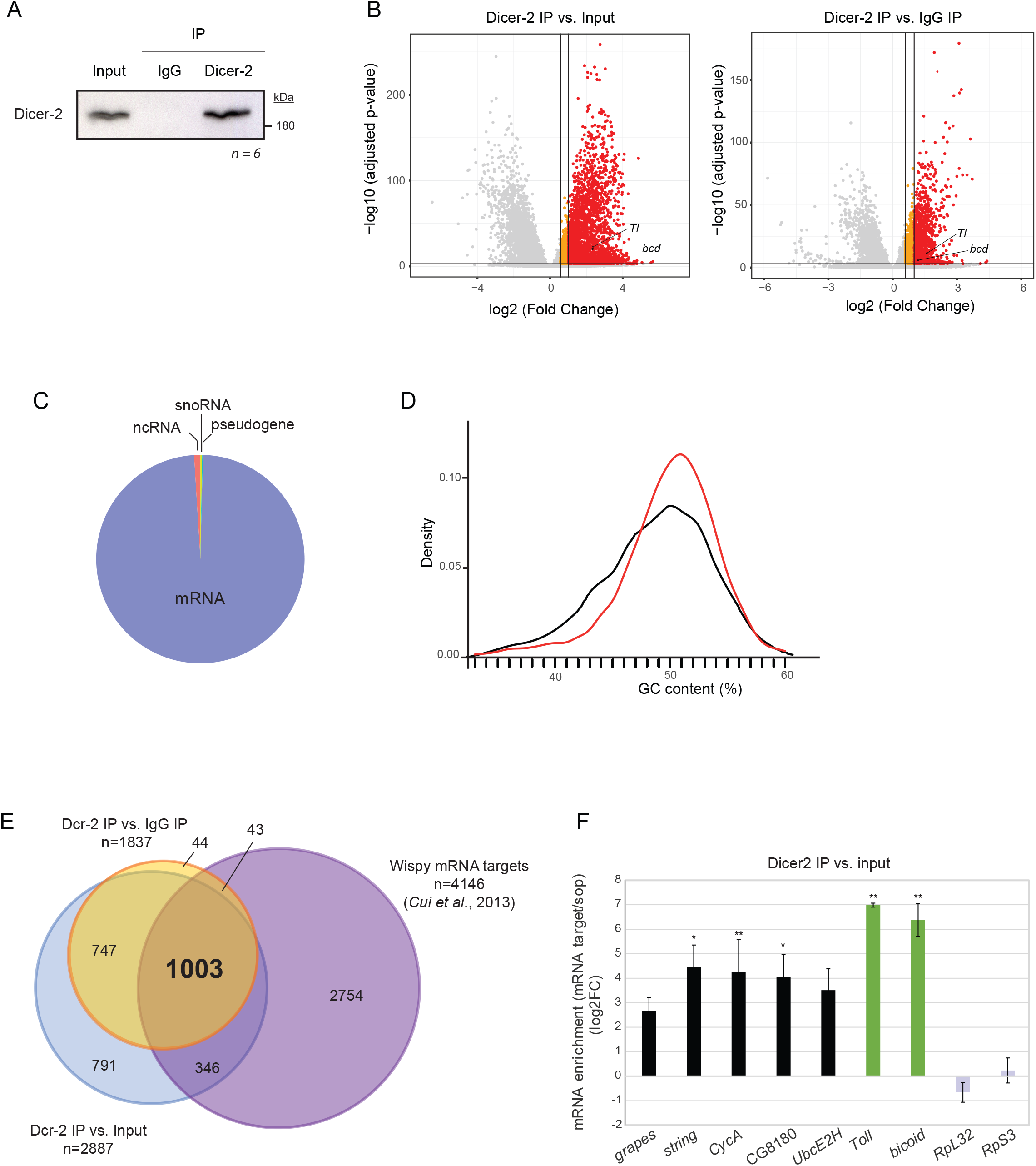
Identification of Dicer-2 mRNA targets. A. Representative image of Dicer-2 immunoprecipitation. IgG, negative control. B. Volcano plots showing RNA enrichment in Dicer-2 IPs with respect to input (*left*) or IgG (*right*). Red, fold enrichment > 2; orange, fold enrichment between 1.5 and 2; grey, not enriched. *Toll* (*Tl*) and *bicoid* (*bcd*) mRNAs are indicated. C. RNA types bound by Dicer-2. D. GC-content of Dicer-2 targets (red) compared to non-targets (black). E. Overlap of Dcr-2 and Wispy mRNA targets.F. Validation of RIP-seq by RT-qPCR. Selected targets of the Dcr-2/Wispy overlap were chosen. The enrichment was normalized to *sop* (*RpS2*), a ribosomal protein mRNA which does not undergo cytoplasmic polyadenylation. *Toll* and *bicoid* mRNAs (green) were used as positive controls, while *RpL32* and *RpS3* mRNAs (grey) were used as non-targets.

Transcripts known to be substrates of cytoplasmic polyadenylation (*Toll, bicoid*) were associated to Dicer-2. In order to understand the potential of Dicer-2 as a cytoplasmic polyadenylation factor, we compared the list of Dicer-2 bound transcripts to that of Wispy polyadenylation substrates identified in a previous study (Cui et al., 2013). About one thousand transcripts were present in both lists, indicating that a large fraction of Dicer-2 targets (57%) represent substrates of cytoplasmic polyadenylation (Figure 1E). Independent validation by RIP/RT-qPCR indeed confirmed binding of Dicer-2 to Wispy substrates, while abundant non-substrate mRNAs coding for ribosomal proteins were not bound (Figure 1F). These results suggest widespread roles of Dicer-2 in cytoplasmic polyadenylation.

### The Dicer-2 protein interactome

In order to identify the Dicer-2 protein interactome of early embryos, we performed co-immunoprecipitation (co-IP) using affinity purified αDicer-2 antibodies. As negative controls, we used either Dicer-2 null embryos or IP with non-specific IgG, each comparison in biological triplicates. Because Dicer-2 null males are sterile, we obtained null embryo extracts by crossing homozygous Dicer-2 null females (*dicer-2*^*L811fsX*^) with wild-type males (*w*^*1118*^), whose contribution to the embryo protein content is only noticeable after 2 hours of development once the maternal-to-zygotic transition has occurred (Figure 2A). Therefore, the male contribution to our early (90’) embryo extracts is negligible, as corroborated by Western blot (Figure 2B). We included treatment with RNase I to identify RNA-independent interactions. As expected, we observed co-immunoprecipitation of Dicer-2 with Wispy in an RNA-independent fashion, and no background in the null or IgG controls (Figures 2B-C). Proteins present in the immunoprecipitates were then identified by tandem mass spectrometry (LC-MSMS).

**Figure 2:**
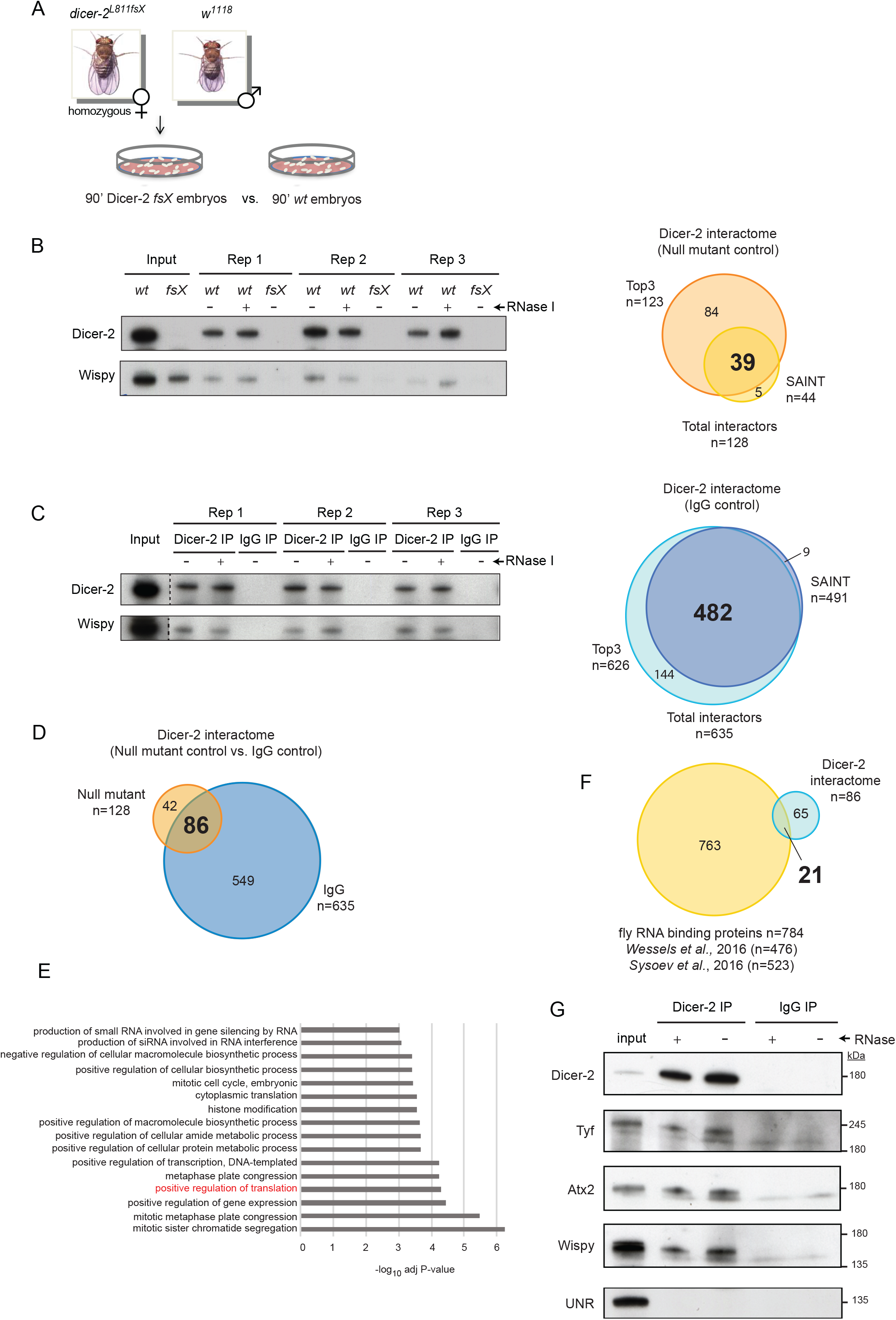
Identification of the Dicer-2 protein interactome. A. Schematic representation of the strategy to obtain Dicer-2 null embryos. B, C. Comparison of wild type (*w*^*1118*^, *wt*) and Dicer-2 null (*fsX*) embryos (B), or αDicer-2 versus IgG IPs (C). In both cases, an immunoblot analysis of three replicates after treatment or not with RNase I is shown on the *left*, and the overlap between SAINT and Top3 analyses is shown on the *right*. D. The Dicer-2 high confidence interactome, defined as the overlap between proteins identified in B and C (86 proteins). E. Gene Ontology analysis of the Dicer-2 high-confidence interactome using Flyenrichr under the term ‘Biological Process’. F. Comparison between the Dicer-2 high-confidence interactome and the *Drosophila* RBPomes identified in 0-2h embryos (Wessels et al. 2016) and across the maternal-to-zygotic transition (Sysoev et al. 2016). G. Validation of Dicer-2 partners by co-immunoprecipitation. Dicer-2, Tyf, Atx2 and Wispy interact in an RNA-independent fashion. UNR was used as a negative control.

Two different bioinformatic procedures were used to identify true Dicer-2 interactors, SAINTexpress and Top3 (Teo et al., 2014; Silva et al., 2006). SAINT (Significant Analysis of INTeractome) uses the spectral counts of all peptides in a protein, while Top3 utilizes the average peak areas of the three most intense peptides per protein. SAINT scored interactions for 854 proteins, of which 44 were found significant (Bayesian FDR ≤ 0.05) over the null control, while Top3 analysis revealed 123 significant interactors. Altogether, 128 proteins were found to interact with Dicer-2 over the null control, of which 39 were found using both SAINT and Top3 (Figure 2B, right and Table S2).

The analyses described above were also applied to find significant Dicer-2 interactors over the IgG control. We found a total of 635 interactors, of which 482 were both SAINT and Top3 hits (Figure 2C, right and Table S2). Overall, 86 proteins showed enrichment in the Dicer-2 IP against both the null and the IgG control, and were categorized as the high-confidence Dicer-2 interactome (Figure 2D). Of these, 70 proteins (81%) were found to interact with Dicer-2 in an RNA-independent manner in all comparisons (Table S2). Therefore, the vast majority of interactions in the Dicer-2 high-confidence set are direct.

We next compared the Dicer-2 high confidence interactome with siRNA-associated factors previously identified by affinity chromatography against two different siRNAs in *Drosophila* embryo extracts (Gerbasi et al., 2010). Only two RNAi factors, R2D2 and Dicer-1, were found to overlap with our list, suggesting pervasive functions of Dicer-2 outside the RNAi machinery. Indeed, GO analysis indicated prevalence of cell division and biosynthetic processes, including translation regulation, among the terms most significantly associated with this group (Figure 2E).

To narrow-down the partners of Dicer-2 potentially involved in cytoplasmic polyadenylation, we searched for Dicer-2-associated RNA binding proteins (RBPs). We compared the Dicer-2 interactome with two recently published mRNA-bound proteomes of fly embryos (Wessels et al., 2016; Sysoev et al., 2016), revealing 21 RBPs as potential co-factors of Dicer-2 (Figure 2F, Table S2). Among these, we focused on RBPs involved in translational control. Notably, in addition to the translation regulators oskar, Fmr1, mushashi, Rasputin and Wispy, we found two proteins, Twenty-four (Tyf) and Ataxin-2 (Atx2), previously shown to stimulate the synthesis of the clock protein Period in a manner that requires Poly(A) binding protein (Lim et al., 2011; Lim and Allada, 2013; Zhang et al., 2013). Furthermore, both the yeast Atx2 homolog Pbp1p and human ATXN2 have been shown to be involved in regulation of poly(A) tail length (Mangus et al., 1998, 2004; Inagaki et al., 2020). Thus, we hypothesized that Dicer-2, Atx2, Tyf and Wispy may form an RNA-independent complex that works in cytoplasmic polyadenylation. Indeed, co-IP experiments corroborated the presence of such complex (Figure 2G).

### Twenty-four, Ataxin-2 and Dicer-2 work in cytoplasmic polyadenylation

To test whether Tyf and Atx2 function in cytoplasmic polyadenylation, we depleted them from embryos by expressing UAS-RNAi constructs under the control of the maternal alphaTub67C driver (Figure 3A). In addition, the capacity of Dicer-2 to polyadenylate targets other than *Toll* mRNA was assessed using Dicer-2 null embryos. Extracts were obtained from these embryos, and polyadenylation efficiencies assessed using poly(A)-test (PAT) assays (Sallés and Strickland, 1995) in comparison wild type embryos. Figure 3A shows that Tyf, Atx2 and Dicer-2 were efficiently depleted. Evaluating five novel Dicer-2 targets indicated that cytoplasmic polyadenylation of four of them (grapes, CG8180, string and UbcE2H) was reduced upon depletion of Atx2 or Tyf, or upon deletion of Dicer-2, while polyadenylation of CycA was not affected by anyone factor (Figure 3B, see quantification on the right). These results indicate that Atx2, Tyf and Dicer-2 act in concert for cytoplasmic polyadenylation of Dicer-2 targets. The results also show that the functions of Atx2 in cytoplasmic polyadenylation are highly conserved, and establish a new role for Tyf in this process, increasing the diversity of RBPs that functionally interact with cytoplasmic poly(A) polymerases (reviewed in Liudkovska and Dziembowski, 2021).

**Figure 3:**
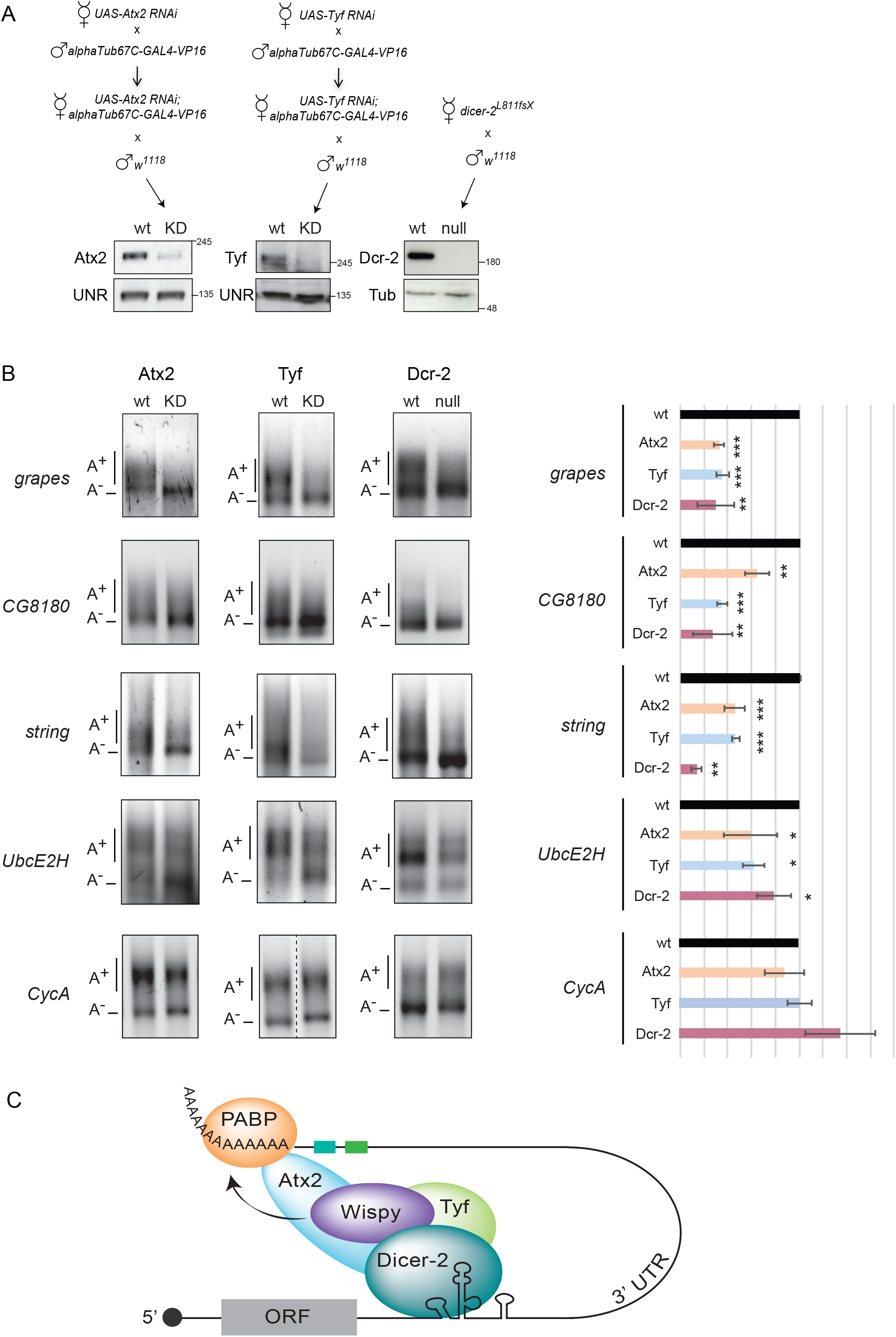
Atx2 and Tyf are involved in cytoplasmic polyadenylation of Dicer-2 targets. A. Schematic representation of the crosses performed to obtain depleted/null extracts (*top*) and efficiency of depletion (*bottom*). B. Efficiency of polyadenylation of Dicer-2 targets in wild type (*w*^*1118*^) versus Atx2-depleted, Tyf-depleted or Dicer-2 null embryos. Quantification of 3 independent experiments is shown on the right. Error bars represent standard deviation. Significance was assessed using Student’s T-test. C. Model for the embryonic non-canonical cytoplasmic polyadenylation complex.

Atx2 has been shown to form mutually exclusive translation activation or repression complexes to regulate circadian clock periodicity and amplitude (Lee et al., 2017). The activation complex contains Lsm12 and Tyf, while the repression complex contains NOT1 and Me31B/DDX6, two proteins involved in deadenylation and miRNA-dependent silencing (Temme et al., 2014; Ostareck et al., 2014). Remarkably, Lsm12 is part of the Dicer-2 high confidence interactome, while NOT1 and Me31B are not (Table S2). These results support a model where Dicer-2 selectively interacts with the Atx2 activation complex to promote cytoplasmic polyadenylation and translation in early embryos (Figure 3C). In sum, our results reveal the composition of a non-canonical cytoplasmic polyadenylation machinery that recruits Wispy for activation of maternal mRNAs, illustrating the growing diversity and plasticity of poly(A) tail length control.

## MATERIALS AND METHODS

### Fly strains and generation of depleted and null embryos

*OregonR and w*^*1118*^ were used as wild-type flies. The following mutant or transgenic stocks were used: *w; Dcr2*^*L811fsX*^*/CyO, amos*^*Roi-1*^ (kindly provided by Dr. Martine Simonelig), *w; UAS-Atx2-RNAi* (Vienna Drosophila Resource Center, VDRC #34955), *w; UAS-Tyf-RNAi* (VDRC #21965) and *w; alphaTub67C-GAL4-VP16; alphaTub67C-GAL4-VP16* (containing the driver in both chromosomes II and III, provided by Dr. Jerome Solon). Flies were maintained on standard food at 25°C.

*Dicer-2* null embryos were obtained by crossing *w; Dcr2*^*L811fsX*^ homozygous females with *w*^*1118*^ males. For the generation of Atx2-depleted embryos, homozygous virgin *w; UAS-Atx2-RNAi* females were crossed with homozygous *alphaTub67C-GAL4-VP16* males, and resulting females were then crossed with *w*^*1118*^ males. A similar strategy was followed to obtain Tyf depleted embryos using *w; UAS-Tyf-RNAi* females.

### Embryo extracts

For Dicer-2 interactome and RIP-Seq analysis, large scale collections of staged *OregonR* 90 min embryos were obtained and processed as described in Coll et al. (2010). For PAT assays, the protocol was scaled down as follows, carrying depleted/null and wild type embryos in parallel. Egg-laying trays were exchanged three times for embryo synchronization, and 90 min embryos collected thereafter. In the case of Dcr-2 null embryos, because of their scarcity, multiple 90 min collections were taken in a period of 6-8 h, kept on ice and pooled. Embryos were washed twice in EW buffer (0.7% NaCl, 0.04% TritonX-100), dechorionated in bleach (1:3 in EW buffer) for 2-3 min, and washed several times with milliQ water. Dechorionated embryos were then washed once with cold DEI buffer (10 mM HEPES pH 7.4, 5 mM DTT and 1x complete protease inhibitor from Roche) followed by homogenization in 1-2 embryo volumes of DEI buffer on ice. The homogenate was centrifuged at 10,000g for 10 min at 4°C. The intermediate cytosolic fraction was carefully collected, avoiding the upper lipid layer and the pellet. Glycerol was added to 10%, and extracts aliquoted, snap frozen and stored at −80°C.

### Antibodies

Rabbit αDicer-2 polyclonal antibodies were raised in house against the first 151 aa of the protein (Coll et al., 2018), and affinity-purified using HiTrap NHS-activated HP columns (GE Healthcare) coupled with His-Dicer-2(1-257)-MBP. αWispy antibodies were kindly provided by Mariana Wolfner (Cui et al., 2008). αTyf and αAtx2 antibodies were provided by Joonho Choe (Lim et al., 2011) and Mani Ramaswamy (McCann et al., 2011), respectively. αUNR antibodies have been described previously (Abaza et al., 2006).

Antibody dilutions for Western blots were as follows: αDicer-2 (1:1000), αTyf (1:500), αAtx2 (1:500), αWispy (1:2000), αUNR (1:1000).

### RNA immunoprecipitation followed by sequencing (RIP-Seq)

Protein-A Dynabeads (Invitrogen) containing affinity purified αDicer-2 antibody to their maximum binding capacity (2.4 µg antibody per 10 µl beads) were prepared by washing beads with 10 volumes of IPP 500 buffer (20 mM HEPES, 500 mM NaCl, 1.5 mM MgCl_2_, 0.5 mM DTT and 0.05 % NP-40) followed by incubation with αDicer-2 antibody for 30 min at room temperature. Antibody-bound beads were washed twice with 10 vol of IPP 500 and IPP containing 150 mM NaCl (IPP 150). A similar procedure was followed to obtain beads with IgG used as negative control.

Embryo extracts were pre-cleared with empty beads equilibrated in IPP 150 for 30 min at 4°C in the presence of 1x complete protease inhibitor cocktail (Roche). Pre-cleared extracts (300-500 µg) were then added to 60 µl of antibody-bound beads in a final volume of 780 µl of IPP 150, and incubated at 4°C for 4h on a rotating wheel. Following IP, beads were pelleted, washed four times with 5 vol of IPP 150, and RNA extracted using TRIZOL. RNA quality was monitored in a Bioanalyzer. An aliquot of the IP was reserved for protein extraction and IP quality control.

Illumina True-seq libraries were prepared at the CRG Genomics facility using the Ribo-zero kit (Illumina) and subjected to 50 bp single-end sequencing. Reads were aligned to the *Drosophila melanogaster* ENSEMBL genome release 87 (BDGP release 6_ISO1 MT/dm6 assembly) using the STAR mapper (version 2.5.2a) (Dobin et al., 2013). Qualimap (version 2.2.1) was used to check the quality of the mapped bam files (García-Alcalde et al., 2012). A raw count of reads per gene was obtained with HTSeq (htseq-count function, version 0.6.1p1) (Anders et al., 2014). The R/Bioconductor package DESeq2 was used to assess differential enrichment between experimental samples. Prior to DESeq2, genes for which the sum of raw counts across all samples was less than 2 were discarded.

### Dicer-2 protein interactome identification

Dicer-2 IPs were performed as indicated above, using 300 µg of wild type (*w*^*1118*^) or Dicer-2 null extracts, keeping the proportions of extract to beads and volumes. Following IP and washes, one half of the Dicer-2 beads were incubated with 100 U RNase I (Ambion) in 100 µl IPP 150 at 25°C for 15 min on a thermomixer. Untreated samples were incubated in parallel without addition of RNase I. Beads were then washed four times with 5 vol of IPP 150 before elution of proteins for mass spectrometry and western blot. For mass spectrometry, samples on beads were washed thrice with 500 µl of 200 mM ammonium bicarbonate (ABC) and resuspended in 60 µl of 6 M urea in 200 mM ABC. Proteins were then reduced by adding DTT (10 µl DTT 10 mM, 37ºC, 60 min), alkylated with iodoacetamide (10 µl of IAM 20 mM, 25ºC, 30 min), diluted with 200 mM ABC to reach a final urea concentration of 1M, and digested overnight with trypsin (1 µg, 37ºC). The peptide mixture was collected and acidified to a final concentration of 5 % formic acid. Samples were desalted using a C18 column, evaporated to dryness and diluted to 10 µl with 0.1 % formic acid in milliQ water. Forty five percent of the peptide mixture was analyzed by LC-MSMS using a 1-hour gradient in the LTQ-Orbitrap Velos Pro mass spectrometer (Thermo Fisher Scientific, San Jose, CA). The data were acquired with Xcalibur software v2.2. and analyzed using the Proteome Discoverer software suite (v1.4.1.14, Thermo Fisher Scientific). The search engine Mascot (v2.5.1 Matrix Science; Perkins et al., 1999) was used for peptide identification. Peptide data was searched against the Uniprot (*UP_Drosophila)* protein database and the identified peptides were filtered to a threshold of 5% FDR.

Protein-protein interactions were assessed using SAINTexpress (Teo et al., 2014) and Top3 (Silva et al., 2006) analysis. SAINT interactors with Bayesian false discovery rate (BFDR) of < 0.05 were included in the Dicer-2 interactome. For Top3 analysis, the log_2_ of the average area of the three most intense peptides of each protein was calculated by Proteome Discoverer. Top3 analysis was constrained by very low abundance or absence of several proteins in the null IP, as well as by missing peak area values for one out of three null mutant replicates. These technical caveats were overcome by semi-quantitative scoring of significant peptide occurrences in wild-type vs null replicates, and by imputation that substitutes the missing values for the mean value of other replicates, respectively. A Student’s t-test was performed between the 3 replicates of each sample condition to identify differentially abundant proteins.

### Small scale immunoprecipitation

Protein-A dynabeads (80 µl) were equilibrated in wash buffer (10 mM Hepes pH8.0, 8 % Glycerol) and blocked by incubation with 600 µg embryo extract for 1 hour at room temperature. After extensive washing, 10 µg of affinity-purified αDicer-2 antibody or IgG were added in wash buffer followed by incubation for 1 hour at room temperature and five additional washes. Beads were mixed with 600 µg of embryo extract in 200 µl DE buffer (10 mM HEPES, 5 mM DTT), supplemented with 100 µl of buffer containing 20 mM Hepes pH 8, 150 mM NaCl, 0.1% NP-40, 1 mM EDTA, 0.5 mM DTT and 1x protein inhibitor cocktail (Roche), and incubated at 25ºC for 1 h on a rotating wheel. Beads were then washed three times with 5 volumes of 1xNET (50 mM Tris-HCl pH7.5, 150 mM NaCl, 0.1% NP-40, 1 mM EDTA). One half of the beads were incubated with 100 U RNase I (Ambion) and 10 µg of RNase A (Sigma) in 40 µl wash buffer with protein inhibitors at room temperature for 20 min. Untreated samples were incubated in parallel without adding RNases. Beads were washed three times with 10 volumes of 1xNET and proteins eluted in SDS buffer for western blot.

### PAT assays

Pools of 100-200 embryos were collected in eppendorf tubes and disrupted with Pellet mixer Cordless (Avantor) in 200 µl Trizol. RNA was obtained using a Maxwell 16 LEV simplyRNA kit (Promega). One microgram of total RNA was used for PAT assays as described in Sallés and Strickland (1995). Gene-specific forward oligonucleotides used for PAT assays were as follows: String (5’-CCGAAACGCAAATGCAAAC-3’), Grapes (5’-CATTGCATTGTTTACGAGTACG-3’), CycA (5’-GCCACCGGACACAACATATTA-3’), UbcE2H (5’-TGTCTGTCGTCGTCATTACGC-3’), CG8180 (5’-CAGCAGCATTAACTGACGAATCGAC-3’).

### RT-qPCR

RNA was reverse-transcribed with SuperScript II (Invitrogen) using a mix of random primers and oligo(dT). The cDNAs were subjected to quantitative real-time PCR using SYBR Green PCR master mix (Applied Biosystems) and ViiA 7 Real-Time PCR System (Applied Biosystems) following the manufacturer’s instructions. Oligonucleotides used in qPCR are as follows: Toll-f (5’-CGCTGCCTTCGCGTCTGTTTGC-3’), Toll-r (5’-GTGTGGAATGCTCGAATAAGTCA-3’), Bicoid-f (5’-ATTGCAATCTGTTAGGCCTCAAG-3’), Bicoid-r (5’-CGGGATCCCGAGTAGAGTAGTTCTTATATATT-3’), Sop-f (5’-CCGTGGTACTGGCATTGTCT-3’), Sop-r (5’-CCGAGTATGCCTGGTAAGGA-3’), Grapes-f (5’-TGAGGAGAATGACCCGATTC-3’), Grapes-r (5’-AACCACCCAGTCTTTCGATG-3’), String-f (5’-CACAAGCGCAACATCATTATC-3’), String-r (5’-CCGGATAGGCGTTGGTATT-3’), CycA-f (5’-GCTGGAGGAGATCACGACTT-3’), CycA-r (5’-TTGTACTTTTCCCGCATGG-3’), CG8180-f (5’-AAGGTCCCAACATCTCTCTGA-3’), CG8180-r (5’-GCTGGGGATGGTATCTCGTA-3’), UbcE2H-f (5’-CAACGATCGCAACAGATCAT-3’), UbcE2H-r (5’-TTTTGCTCTTCCATCCGTTC-3’), RpL32-f (5’-TGCCCACCGGATTCAAGA-3’), RpL32-r (5’-AAACGCGGTTCTGCATGAG-3’), RpS3-f (5’-CGATTTCCAAGAAACGCAAG-3’), RpS3-r (5’-CGAGTCAGGAACTCGTTCAA-3’).

## ACKNOWLEDGEMENTS

We thank Mariana Wolfner, Joonho Choe and Mani Ramaswamy for kindly sharing their antibodies. We also thank the CRG Proteomics, Genomics, Bioinformatics and Protein Technologies Facilities for protein identification, RNA sequencing, data analysis and αDicer-2 affinity purification, respectively. H-P. N. was supported by a La Caixa fellowship. This work was supported by grants from the Spanish Ministry of Science and Innovation (PGC2018-099697-B-I00 and BFU2015-68741), the Catalan Agency for Research and Universities (2017SGR534) and the Centre of Excellence Severo Ochoa.

## AUTHOR CONTRIBUTIONS

HP.N. and F.G. designed the study. HP.N., T.G. and O.C. performed the experiments. F.G. conceived and supervised the project. All authors contributed to writing and editing the manuscript.

## REFERENCES

Abaza I, Coll O, Patalano S, Gebauer F (2006) Drosophila UNR is required for translational repression of male-specific lethal 2 mRNA during regulation of X-chromosome dosage compensation. Genes Dev 20:380–389.

Anders S, Pyl PT, Huber W (2015) HTSeq--a Python framework to work with high-throughput sequencing data. Bioinformatics 31: 166–169

Charlesworth A, Meijer HA, de Moor CH (2013) Specificity factors in cytoplasmic polyadenylation. Wiley Interdiscip Rev RNA 4: 437–461

Coll O, Guitart T, Villalba A, Papin C, Simonelig M, Gebauer F (2018) Dicer-2 promotes mRNA activation through cytoplasmic polyadenylation. RNA 24: 529–539

Coll O, Villalba A, Bussotti G, Notredame C, Gebauer F (2010) A novel, noncanonical mechanism of cytoplasmic polyadenylation operates in Drosophila embryogenesis. Genes Dev 24: 129–134

Cui J, Sackton KL, Horner VL, Kumar KE, Wolfner MF (2008) Wispy, the Drosophila homolog of GLD-2, is required during oogenesis and egg activation. Genetics 178: 2017–2029

Cui J, Sartain CV, Pleiss JA, Wolfner MF (2013) Cytoplasmic polyadenylation is a major mRNA regulator during oogenesis and egg activation in Drosophila. Dev Biol 383: 121–131

D’Ambrogio A, Nagaoka K, Richter JD (2013) Translational control of cell growth and malignancy by the CPEBs. Nat Rev Cancer 13: 283–290

Dobin A, Davis CA, Schlesinger F, Drenkow J, Zaleski C, Jha S, Batut P, Chaisson M, Gingeras TR (2013) STAR: ultrafast universal RNA-seq aligner. Bioinformatics 29: 15–21

Dufourt J, Bontonou G, Chartier A, Jahan C, Meunier AC, Pierson S, Harrison PF, Papin C, Beilharz TH, Simonelig M (2017) piRNAs and Aubergine cooperate with Wispy poly(A) polymerase to stabilize mRNAs in the germ plasm. Nat Commun 8: 1305

Fernandez-Miranda G, Mendez R (2012) The CPEB-family of proteins, translational control in senescence and cancer. Ageing Res Rev 11: 460–472

Garcia-Alcalde F, Okonechnikov K, Carbonell J, Cruz LM, Gotz S, Tarazona S, Dopazo J, Meyer TF, Conesa A (2012) Qualimap: evaluating next-generation sequencing alignment data. Bioinformatics 28: 2678–2679

Gerbasi VR, Golden DE, Hurtado SB, Sontheimer EJ (2010) Proteomics identification of Drosophila small interfering RNA-associated factors. Mol Cell Proteomics 9: 1866–1872

Gewartowska O, Aranaz-Novaliches G, Krawczyk PS, Mroczek S, Kusio-Kobialka M, Tarkowski B, Spoutil F, Benada O, Kofronova O, Szwedziak P et al (2021) Cytoplasmic polyadenylation by TENT5A is required for proper bone formation. Cell Rep 35: 109015

Grabek KR, Diniz Behn C, Barsh GS, Hesselberth JR, Martin SL (2015) Enhanced stability and polyadenylation of select mRNAs support rapid thermogenesis in the brown fat of a hibernator. Elife 4

Inagaki H, Hosoda N, Tsuiji H, Hoshino SI (2020) Direct evidence that Ataxin-2 is a translational activator mediating cytoplasmic polyadenylation. J Biol Chem 295: 15810–15825

Ivshina M, Lasko P, Richter JD (2014) Cytoplasmic polyadenylation element binding proteins in development, health, and disease. Annu Rev Cell Dev Biol 30: 393–415

Kojima S, Sher-Chen EL, Green CB (2012) Circadian control of mRNA polyadenylation dynamics regulates rhythmic protein expression. Genes Dev 26: 2724–2736

Kozlov E, Shidlovskii YV, Gilmutdinov R, Schedl P, Zhukova M (2021) The role of CPEB family proteins in the nervous system function in the norm and pathology. Cell Biosci 11: 64

Lee J, Yoo E, Lee H, Park K, Hur JH, Lim C (2017) LSM12 and ME31B/DDX6 Define Distinct Modes of Posttranscriptional Regulation by ATAXIN-2 Protein Complex in Drosophila Circadian Pacemaker Neurons. Mol Cell 66: 129–140 e127

Lim C, Allada R (2013) ATAXIN-2 activates PERIOD translation to sustain circadian rhythms in Drosophila. Science 340: 875–879

Lim C, Lee J, Choi C, Kilman VL, Kim J, Park SM, Jang SK, Allada R, Choe J (2011) The novel gene twenty-four defines a critical translational step in the Drosophila clock. Nature 470: 399–403

Liudkovska V, Dziembowski A (2021) Functions and mechanisms of RNA tailing by metazoan terminal nucleotidyltransferases. Wiley Interdiscip Rev RNA 12: e1622

Mangus DA, Amrani N, Jacobson A (1998) Pbp1p, a factor interacting with Saccharomyces cerevisiae poly(A)-binding protein, regulates polyadenylation. Mol Cell Biol 18: 7383–7396

Mangus DA, Smith MM, McSweeney JM, Jacobson A (2004) Identification of factors regulating poly(A) tail synthesis and maturation. Mol Cell Biol 24: 4196–4206

McCann C, Holohan EE, Das S, Dervan A, Larkin A, Lee JA, Rodrigues V, Parker R, Ramaswami M (2011) The Ataxin-2 protein is required for microRNA function and synapse-specific long-term olfactory habituation. Proc Natl Acad Sci U S A 108: E655–662

Norvell A, Wong J, Randolph K, Thompson L (2015) Wispy and Orb cooperate in the cytoplasmic polyadenylation of localized gurken mRNA. Dev Dyn 244: 1276–1285

Ostareck DH, Naarmann-de Vries IS, Ostareck-Lederer A (2014) DDX6 and its orthologs as modulators of cellular and viral RNA expression. Wiley Interdiscip Rev RNA 5: 659–678

Passmore LA, Coller J (2021) Roles of mRNA poly(A) tails in regulation of eukaryotic gene expression. Nat Rev Mol Cell Biol

Perkins DN, Pappin DJ, Creasy DM, Cottrell JS (1999) Probability-based protein identification by searching sequence databases using mass spectrometry data. Electrophoresis 20: 3551–3567

Pique M, Lopez JM, Foissac S, Guigo R, Mendez R (2008) A combinatorial code for CPE-mediated translational control. Cell 132: 434–448

Rybak-Wolf A, Jens M, Murakawa Y, Herzog M, Landthaler M, Rajewsky N (2014) A variety of dicer substrates in human and C. elegans. Cell 159: 1153–1167

Salles FJ, Strickland S (1995) Rapid and sensitive analysis of mRNA polyadenylation states by PCR. PCR Methods Appl 4: 317–321

Silva JC, Gorenstein MV, Li GZ, Vissers JP, Geromanos SJ (2006) Absolute quantification of proteins by LCMSE: a virtue of parallel MS acquisition. Mol Cell Proteomics 5: 144–156

Sysoev VO, Fischer B, Frese CK, Gupta I, Krijgsveld J, Hentze MW, Castello A, Ephrussi A (2016) Global changes of the RNA-bound proteome during the maternal-to-zygotic transition in Drosophila. Nat Commun 7: 12128

Temme C, Simonelig M, Wahle E (2014) Deadenylation of mRNA by the CCR4-NOT complex in Drosophila: molecular and developmental aspects. Front Genet 5: 143

Teo G, Liu G, Zhang J, Nesvizhskii AI, Gingras AC, Choi H (2014) SAINTexpress: improvements and additional features in Significance Analysis of INTeractome software. J Proteomics 100: 37–43

Villalba A, Coll O, Gebauer F (2011) Cytoplasmic polyadenylation and translational control. Curr Opin Genet Dev 21: 452–457

Wessels HH, Imami K, Baltz AG, Kolinski M, Beldovskaya A, Selbach M, Small S, Ohler U, Landthaler M (2016) The mRNA-bound proteome of the early fly embryo. Genome Res 26: 1000–1009

Zhang Y, Ling J, Yuan C, Dubruille R, Emery P (2013) A role for Drosophila ATX2 in activation of PER translation and circadian behavior. Science 340: 879–882

